# OLIGONUCLEOTIDES TARGETING THE 3’ SPLICE SITE DOWNSTREAM OF A MICROEXON AS AN INNOVATIVE THERAPY FOR AUTISM

**DOI:** 10.1101/2024.10.25.620162

**Authors:** Ainhoa Martinez-Pizarro, Sara Picó, Lise Lolle Holm, Thomas K Doktor, Brage S Andresen, José J Lucas, Lourdes R Desviat

## Abstract

**Background:** Microexons are highly conserved and mostly neuronal-specific 3-27 nucleotide exons, enriched in genes linked to autism spectrum disorders (ASD). We have previously shown decreased inclusion of a neuronal specific 24 bp microexon (exon 4) of the translational regulator *CPEB4* in brains of idiopathic ASD cases and that this leads to CPEB4 aggregation and subsequent under-expression of multiple high confidence ASD-risk genes. Furthermore, enhanced skipping of the *CPEB4* microexon is also a novel etiological mechanism in schizophrenia (SCZ).

**Methods:** In this work we designed and tested in neuroblastoma cells a series of splice switching antisense oligonucleotides (SSO) targeting the *CPEB4* genomic region surrounding the microexon.

**Results:** SSOs targeting candidate intronic regions near the microexon resulted in a decrease in microexon inclusion by blocking hnRNPC/PTPB1 binding, thus mimicking the isoform imbalance observed in ASD. However, based on the kinetic coupling model correlating transcriptional elongation with splicing regulation, we identified SSOs targeting downstream 3’ splice site of exon 5 that favoured microexon inclusion in a dose-dependent manner and resulted in increased protein levels of *FOXP1* and *AUTS2,* two high-confidence ASD risk genes that are known targets of CPEB4 and whose protein levels are reduced in ASD.

**Conclusions:** These results deepen our understanding of the complex splicing regulation of microexons and open new applications of SSOs to treat diseases such as ASD and SCZ that are characterized by altered microexon inclusion.

## BACKGROUND

Autism spectrum disorder (ASD) is a neurodevelopmental and neuropsychiatric disorder characterized by deficits in social interaction and communication, language development, and repetitive behaviours. It affects more than 1% of children and is highly heterogeneous with respect to its presentation and contributing genetic factors. A minority of ASD cases are caused by highly penetrant single-gene mutations or chromosomal abnormalities, while most of the cases are idiopathic [1]. Despite the extensive heterogeneity associated with ASD, common molecular and cellular mechanisms operate in this disorder, such as synaptic function, activity-dependent transcription, and splicing alterations, specifically affecting neuronal microexons [2,3].

Microexons are 3-27 nucleotide long exons that exhibit tissue-specific alternative splicing (AS) and encode for amino acids located in regions with critical roles in protein-protein interactions and function [4]. A major feature of the neuronal microexon alternative splicing (AS) program is that it shows decreased inclusion in brains of patients with ASD, relative to matched controls [3,5]. In this regard, the brains of individuals with idiopathic ASD show decreased inclusion of the neuron-specific 24-nucleotide microexon (exon 4) of cytoplasmic polyadenylation element binding protein 4 (*CPEB4*) gene. CPEBs are RNA binding proteins (RBPs) that recognize transcripts that harbour cytoplasmic polyadenylation element (CPE) sequences (consensus UUUUUAU) in their 3’UTR (representing approximately 40% of the transcriptome) [6,7]. CPEBs repress or activate translation of their target transcripts by inducing, respectively, shortening or elongating their poly(A) tail [8]. It is well established that CPEB-regulated polyadenylation plays a key role in early development, synaptic plasticity and metabolism [8–10]. Alteration of CPEBs and subsequent transcriptome polyadenylation have been associated to different pathologies, notably to central nervous system diseases such as epilepsy [11], ASD [6,12], neurodegenerative disease [13] and schizophrenia (SCZ) [14].

CPEB4-target transcripts are overrepresented among high confidence ASD-risk genes and the imbalance in transcript isoforms with and without microexon 4 (*CPEB4*ex4+/ex4-) in idiopathic ASD brains [6]. Such imbalance leads to abnormal aggregation of CPEB4 [15,16] and correlates with deadenylation of ASD risk genes, with concomitant decreased protein levels [6]. Moreover, mice overexpressing the isoform lacking microexon 4 (TgCPEB4Δ4 mice) recapitulate the deadenylation and decreased protein levels of high confidence ASD risk genes and they also exhibit ASD-like behavioural phenotypes [6]. Recently, a similar pattern was shown in SCZ, with decreased inclusion of *CPEB4* microexon in SCZ probands, correlating with decreased protein levels of CPEB4-target SCZ-associated genes in antipsychotic-free individuals [14]. Mice with mild overexpression of CPEB4Δ4 showed decreased protein levels of CPEB4-target SCZ genes and SCZ-linked behaviours [14].

AS is a highly regulated process influenced by splicing factors and RBPs, acting in *trans,* that bind to *cis* motifs (enhancers or silencers) present on the pre-mRNA, to help or hinder recruitment of the spliceosome [17]. Alternatively spliced microexons display a striking evolutionary conservation and, consistent with this, upstream and downstream flanking intronic sequences are on average more conserved than intron sequences flanking longer alternative exons [3]. This suggests that intronic elements proximal to the microexon splice sites may facilitate their recognition, thus overcoming the steric hindrance for spliceosome assembly resulting from their small length. A number of RBPs that bind to intronic consensus motifs adjacent to microexons and affecting AS of subsets of microexons have recently been identified. Most neural microexons are regulated by the neuronal-specific splicing factor nSR100/SRRM4, which forms a complex with the SR-related proteins, Srsf11 and Rnps1, binding upstream to a bi-partite intronic splicing enhancer element (UCUCUCN1-50UGC) required for microexon recognition and splicing [18]. A small subset of microexons are regulated by other RBPs such as RBFOX, NOVA, QK1 or PTPB1, binding to distinct sites upstream or downstream of the microexon [19]. Additional factors may also influence microexon AS, such as the rate of transcriptional elongation (kinetic coupling mechanism), chromatin structure, and secondary RNA structure that modulates the binding of splice factors to pre-mRNA [20].

Effective splicing modulation has entered the clinic in recent years as a therapeutic approach to pathogenic splicing defects, by use of splice switching antisense oligonucleotides (SSOs) [21–23], with neuronal tissues demonstrated to be amenable for efficient SSO delivery [24]. Following binding to target pre-mRNA according to Watson-Crick pairing rules, SSOs modulate splicing by steric blocking mechanisms, hindering binding of splice factors and RBPs to specific regulatory sequences. Depending on the target sequence, SSOs may promote exon skipping (as exemplified by the SSOs approved for Duchenne muscular dystrophy) [25] or exon inclusion (like Spinraza used in SMA) [26]. Different classes of chemical modifications of the nucleic acid backbone and sugar moiety have been introduced to enhance the nuclease stability and target affinity [23,24].

In this work we employed SSOs targeting different *CPEB4* gene regions with the aim of modulating microexon 4 splicing. We show that SSOs hybridizing to the 3’ splice site of exon 5 favour microexon inclusion and we confirm that SSO treatment in neuroblastoma cells results in an increase in protein products of high-confidence ASD risk genes which are regulated by *CPEB4*. The results provide the proof of concept of the therapeutic potential of this approach for neurological diseases characterized by microexon mis-splicing.

## METHODS

### Cell culture

SH-SY5Y human neuroblastoma and N2A murine neuroblastoma cells were obtained from an in-house collection. Cells were grown in Dulbecco’s Modified Eagle Medium (DMEM, Sigma Aldrich) supplemented with 10% fetal bovine serum (FBS, Gibco), 1% glutamine and 0.1% antibiotic mix (penicillin/streptomycin) under standard cell culture conditions (37°C, 95% relative humidity, 5% CO_2_).

### SSO design

A series of *CPEB4* gene targeting (human and mouse specific sequences) SSOs were designed and synthesized with 2’OMe modification and phosphorothioate (PS) backbones (Supplementary Table 1). Specificity of the selected SSOs was verified using BLAST (https://blast.ncbi.nlm.nih.gov/Blast.cgi). RNA SSOs with the same chemistry were designed, targeting the 3’ splice acceptor site of *TAF1* exon 35 or *eIF4G1* exon 3 (Supplementary Table 1). A scrambled SSO (SCR) not complementary to *CPEB4, eIF4G1* or *TAF1* sequences was used as control (Supplementary Table 1). All SSOs were provided by Integrated DNA Technologies, Leuven, Belgium.

### Minigenes

The *CPEB4* minigene includes from exon 2 to exon 6, with 300 bp of introns dowsntrean and upstream of exons. The minigene was designed, synthesized, and subsequently cloned into the pcDNA3.1+ vector between the restriction enzymes KpnI and XhoI, using the GeneArt service (ThermoFisher Scientific, USA). We generated mutant minigenes containing various deletions and point mutations through site-directed mutagenesis, employing the QuikChange Lightning Kit (Agilent Technologies, Santa Clara, CA, USA). The identity of these constructs was subsequently validated through Sanger sequencing.

### Reverse transfection

Approximately 1×10^6^ cells for SH-SY5Y and 8×10^5^ for N2A were seeded in 6-well plates with 25 pmol (10 nM) 50 pmol (20 nM), 100 pmol (40 nM) of each 2’OMe-PS SSO, or with 2 µg of each minigene, which were reverse transfected into the cells using Lipofectamine 2000 transfection reagent (Invitrogen). Forty-eight hours after transfection cells were harvested for total RNA isolation using Trizol (Invitrogen). Quantification and quality determination of RNA was done on a Nanodrop One (Thermo Scientific).

### RT-PCR

cDNA synthesis was performed using 750 ng of total RNA. Splicing analysis of *CPEB4* transcripts was performed by PCR with primers located in exon 2 and exon 5 or exon 6 (Supplementary Table 2) to amplify the four *CPEB4* splicing isoforms (full-length, Δ4, Δ3, Δ3Δ4). PCR was carried out according to the following protocol: 10 min at 94°C + 33 cycles (30 s at 94°C + 30 s at 58°C + 2 min at 72°C) + 10 min at 72°C.

In the case of minigenes, the primers used are located in exon 3 and between exon 6 and the vector (Exon 6-pcDNA3.1 R). The following protocol was used for PCR amplification: 5 min at 95°C +36 cycles (25 s at 95°C + 25 s at 58°C + 40 s at 72°C) +7 min at 72°C. The endpoint PCR amplification products were subsequently analyzed using 4% agarose gel electrophoresis.

RT-PCR analysis of *TAF1* transcripts was performed using primers located in exon 34 and exon 36 (Supplementary Table 2). Similarly, for *eIF4G1* transcripts, primers in exon 2 and exon 4 were used (Supplementary Table 2). The same PCR protocol described above for minigenes was followed, with a melting temperature of 60°C during the annealing step. PCR products were resolved on a 3% agarose gel. All experiments were conducted in duplicate.

### qRT-PCR

cDNA synthesis was performed using 750 ng of total RNA. Quantification was performed by real-time PCR using a LightCycler® 480 (Roche) in combination with PerfeCta SYBR® Green FastMix (Quanta) and 0.25 μM of primer pair was used (Supplementary Table 2). The mRNA levels were normalized first relative to the GAPDH/Gapdh gene expression and then relative to total RNA in each sample.

### Western blotting

Approximately 8×10^5^ N2A cells were seeded in each well in a 6-well plate (Falcon) with 100 pmol (40 nM) of each 2’OMe-PS SSOs, which was reverse transfected into the cells using Lipofectamine 2000 transfection reagent (Invitrogen). Ninety-six hours after transfection cells were frozen at −20°C and then scrapped in ice-cold extraction buffer (20 mM HEPES pH 7.4, 100 mM NaCl, 20 mM NaF, 1% Triton X-100, 1 mM sodium orthovanadate, 1 μM okadaic acid, 5 mM sodium pyrophosphate, 30 mM β-glycerophosphate, 5 mM EDTA and protease inhibitors (Complete, Roche, Cat. No 11697498001). Between 10 and 20 μg of total protein were electrophoresed on 10% SDS-polyacrylamide gels, transferred to a nitrocellulose blotting membrane (Amersham Protran 0.45 μm, GE Healthcare Life Sciences, 10600002) and blocked in TBS-T (150 mM NaCl, 20 mM Tris–HCl, pH 7.5, 0.1% Tween 20) supplemented with 5% non-fat dry milk. Membranes were incubated overnight at 4°C with the primary antibody FOXP1 (1:1000, Abcam, ab16645) and AUTS2 (1:1000, Sigma, HPA000390) in TBS-T supplemented with 5% non-fat dry milk, washed with TBS-T and next incubated with secondary HRP-conjugated anti-rabbit IgG (1:2000, DAKO, P0448) and developed using the ECL detection kit (PerkinElmer, NEL105001EA). Antibody against Vinculin (1:1000, Abcam, ab129002) was used as loading control. Images were scanned with a densitometer (Bio-Rad, GS-900) and quantified with Image Lab 5.2 (Bio-Rad).

### RNA affinity experiments

A series of 3’-end biotinylated RNA oligonucleotides (LGC Biosearch Technologies; Risskov, Denmark) (Figure 5) covering the *CPEB4* intronic targeted by SSO5 was used to perform affinity purification of A-binding proteins as previously described [27]. 1000 pmol of each 3’ end biotin-coupled RNA oligonucleotide was immobilized in 50 µl streptavidin-coupled magnetic beads; Dynabeads M-280 Streptavidin (Invitrogen; Oslo, Norway), and incubated with RPE-1 nuclear extract (extracted from RPE-1 cells with nuclear extraction kit, abcam, ab113474). Proteins were eluted in XT sample buffer (Bio-Rad Laboratories; Hercules, CA) and analyzed by western blotting with immunodetection using primary antibodies against SRRM4 (1:1000; Aviva, ARP60372_P050), RBFOX2 (1:1000; Nordic biosite, A300-864A), hnRNPC (1:1000; Cell signaling technologies, D6S3N) and PTBP1 (1:1000; Santa cruz, sc-16547), and secondary antibodies anti-rabbit (1:45000, ThermoFisher Scientific, A16104) for the first three and anti-goat (1:10000, Santa Cruz, sc-2056) for PTBP1.

### Data analysis

Statistical analysis was performed with GraphPad Prism version 10.2.3. Data are represented as means ± SEM. For multiple comparisons, data were analyzed by one-way ANOVA. Critical values for significance of \**P*<0.05, \*\**P*<0.01 or \*\*\**P*<0.001 were used throughout the study.

## RESULTS

### Identifying SSOs that modulate splicing of the *CPEB4* microexon

There are four *CPEB4* transcript isoforms present in human brain tissue, resulting from AS of two consecutive exons: the 51-nucleotide exon 3 and the 24-nucleotide neuron-specific microexon 4 [6] (Figure 1A). As mentioned in a previous study, the brains of individuals with idiopathic ASD present an imbalance favoring isoforms lacking exon 4 [6] (Figure 1B). In order to develop an appropriate cellular model as a tool in which to test for splice modulation RNA therapies, which could correct this imbalance by increasing microexon inclusion, we first characterized the *CPEB4* transcript profile in human and murine neuroblastoma cell lines. As shown in Figure 1C, both in N2A and in SH-SY5Y we could clearly detect all four transcript isoforms: full length (FL), without exon 4 (Δ4), without exon 3 (Δ3) and without exons 3,4 (Δ3Δ4). As expected, the isoforms including exon 4 were not detected in non-neuronal cell lines (hepatoma cells, Hep3B and human fibroblasts, CC2509) (Figure 1C).

**Figure 1:**
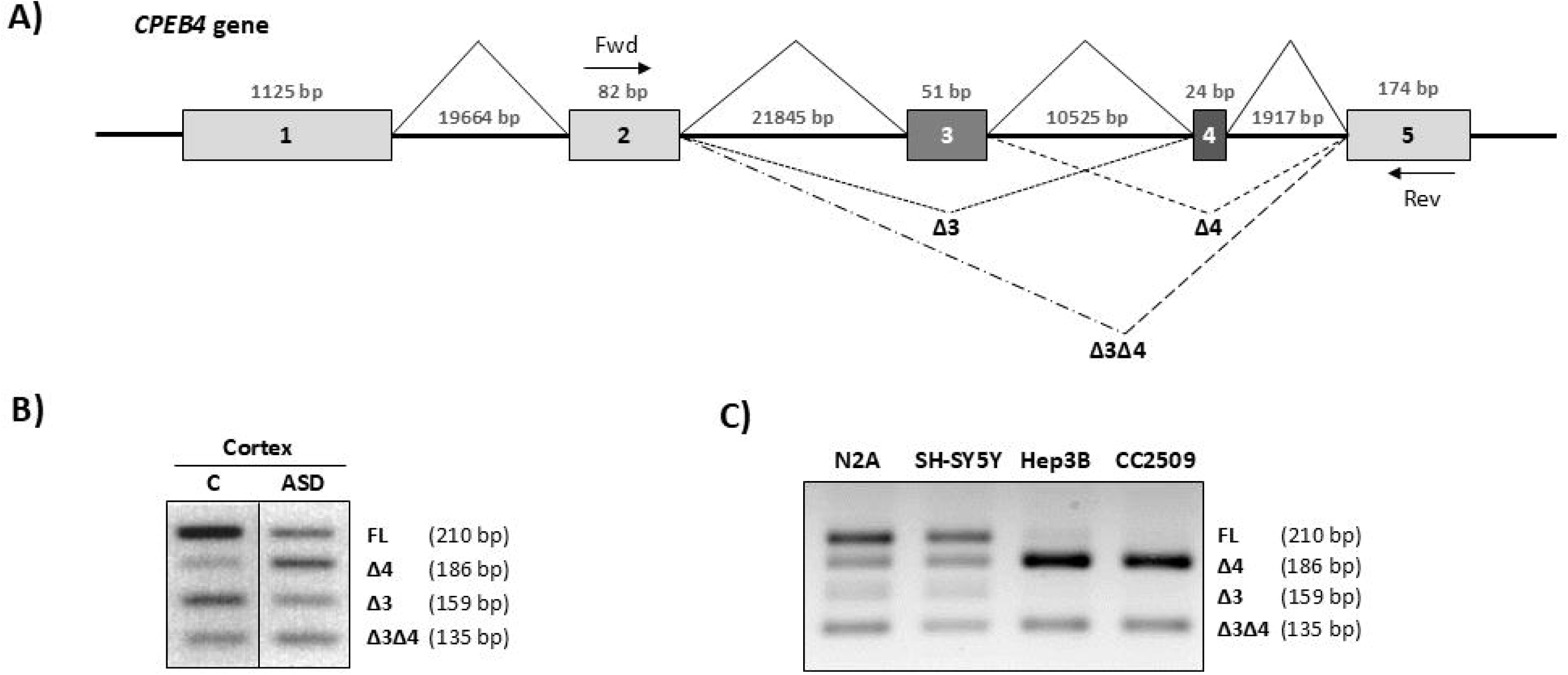
Expression of *CPEB4* transcripts. (A) Schematics of the region of the *CPEB4* gene from exons 1 to 5 indicating AS isoforms and the target sites for forward (Fwd) and reverse (Rev) primers used for amplification; (B) *CPEB4* splice isoforms in control and idiopathic ASD cortex samples; (C) *CPEB4* splice isoforms in control fibroblast (CC2509), Hep3B and SH-SY5Y and N2A neuroblastoma cell lines. Results are representative of two independent experiments. Shown on the right is the identity of the bands and their size. FL, full length; Δ4, without microexon 4; Δ3, without exon 3; Δ3Δ4, without exons 3 and 4.

So far, splicing silencers involved in the regulation of *CPEB4* exon 4 splicing, which could be targeted to promote its inclusion, have not been identified. Initially, we therefore predicted by bioinformatic analysis binding sites for RBP proteins of the hnRNP family, upstream and downstream of exon 4, and designed a series of SSOs targeting these intronic regions (Figure 2A) (Supplementary Table 1). SSOs were transfected in SH-SY5Y cells and RT-PCR analysis was employed to analyze transcript isoforms.

**Figure 2.**
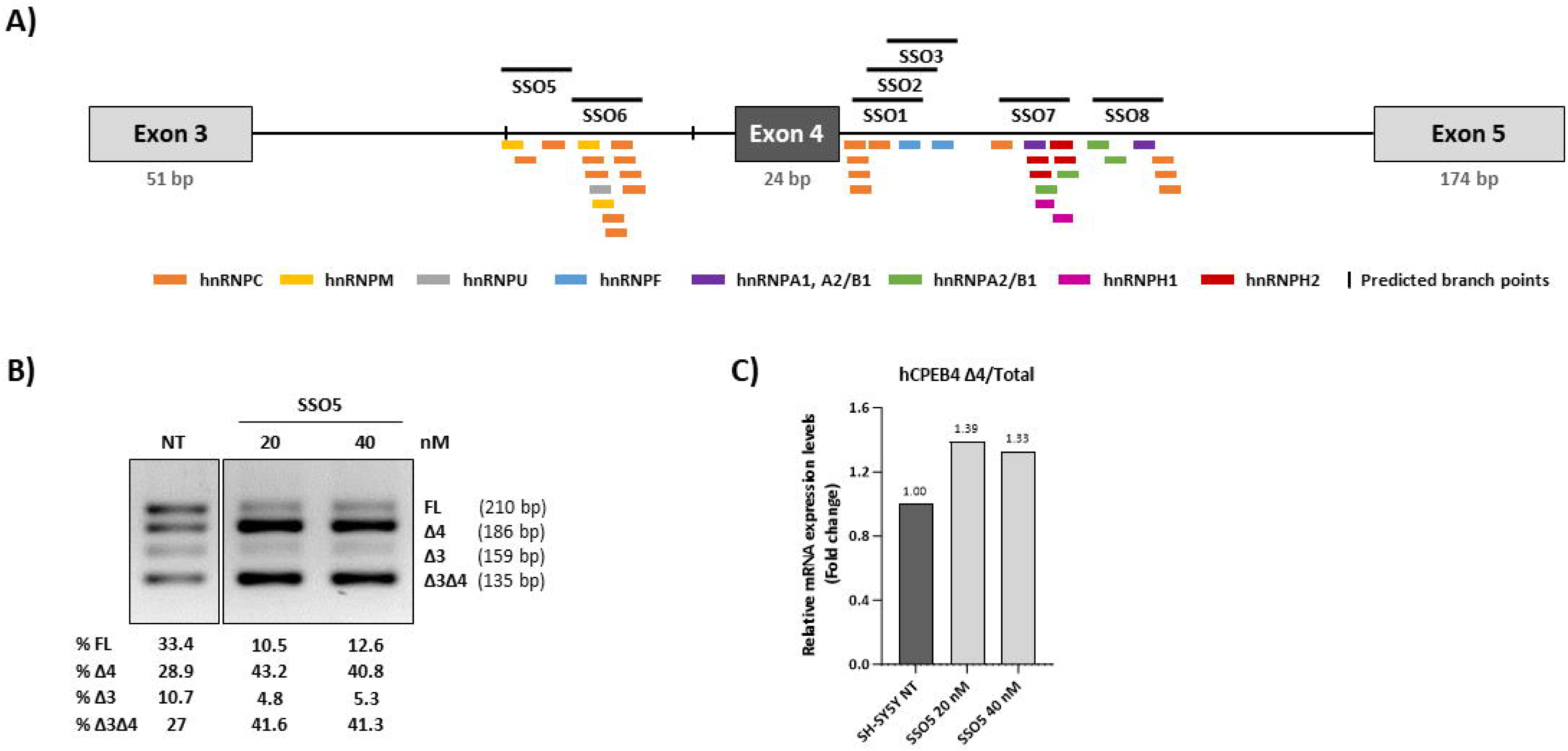
Screening of SSOs targeting intronic regions surrounding exon 4. (A) Schematics of the *CPEB4* gene showing the regions targeted by SSOs 1-8. (B) Gel showing the RT-PCR results after transfection with SSO5 at different concentrations in SH-SY5Y cells. The identity of the bands is shown on the right. These results are representative of two independent experiments. C) RT-qPCR results expressed as fold change in the ratio of isoforms without exon 4 and total (includes all isoforms), relative to the levels in non-transfected cells (NT).

This first screening identified SSO5 as a negative SSO that favored exon 4 exclusion (decrease in transcripts with exon 4 and corresponding increase in Δ4 isoforms) (Figure 2B), while the rest of the SSOs had no significant effect on *CPEB4* transcript profile. The ratio of isoforms without microexon 4 (CPEB4Δex4/total) was increased up to 1,4-fold with SSO5 treatment (Figure 2C), which thus represented a way to generate a cellular model of *CPEB4* mis-splicing, recapitulating the situation characterized in idiopathic ASD brain samples.

As targeting adjacent microexon intronic regions did not reveal any potential therapeutic SSO that could correct ASD associated *CPEB4* mis-splicing, favoring exon 4 inclusion, we decided to use an alternative strategy, namely targeting the 3’ splice acceptor site of exon 5. This approach, which had not been tested before for microexons, is based on the hypothesis that increased upstream exon inclusion will result from lowering of the availability of downstream competing splice sites, generally associated to a slow transcriptional rate (kinetic coupling model) [20]. This mechanism was previously shown to result in increased inclusion of exon 7 in the *SMN2* gene, albeit concomitant with intron 7 retention, thus with no therapeutic benefit for SMA [28,29]. Moreover, we have recently further corroborated the kinetic model by demonstrating that the strength of downstream 3’ splice acceptor sites may determine inclusion of upstream exons like *ACADM* exon 5, *ATM* exon 40 and *BRCA2* exon 3 [30,31]. To test this hypothesis in the *CPEB4* microexon context, we designed overlapping SSOs of different lengths (18-mer to 25-mer) hybridizing to the region encompassing the 3’ splice acceptor site of exon 5 of the *CPEB4* gene, as well as exon 5 sequences and intronic sequences upstream (Figure 3A). The region comprising nucleotides −45 to −5 relative to exon 5 contain predicted branch point (BP) and polypyrimidine sequences, and no SSOs were designed in this region.

**Figure 3:**
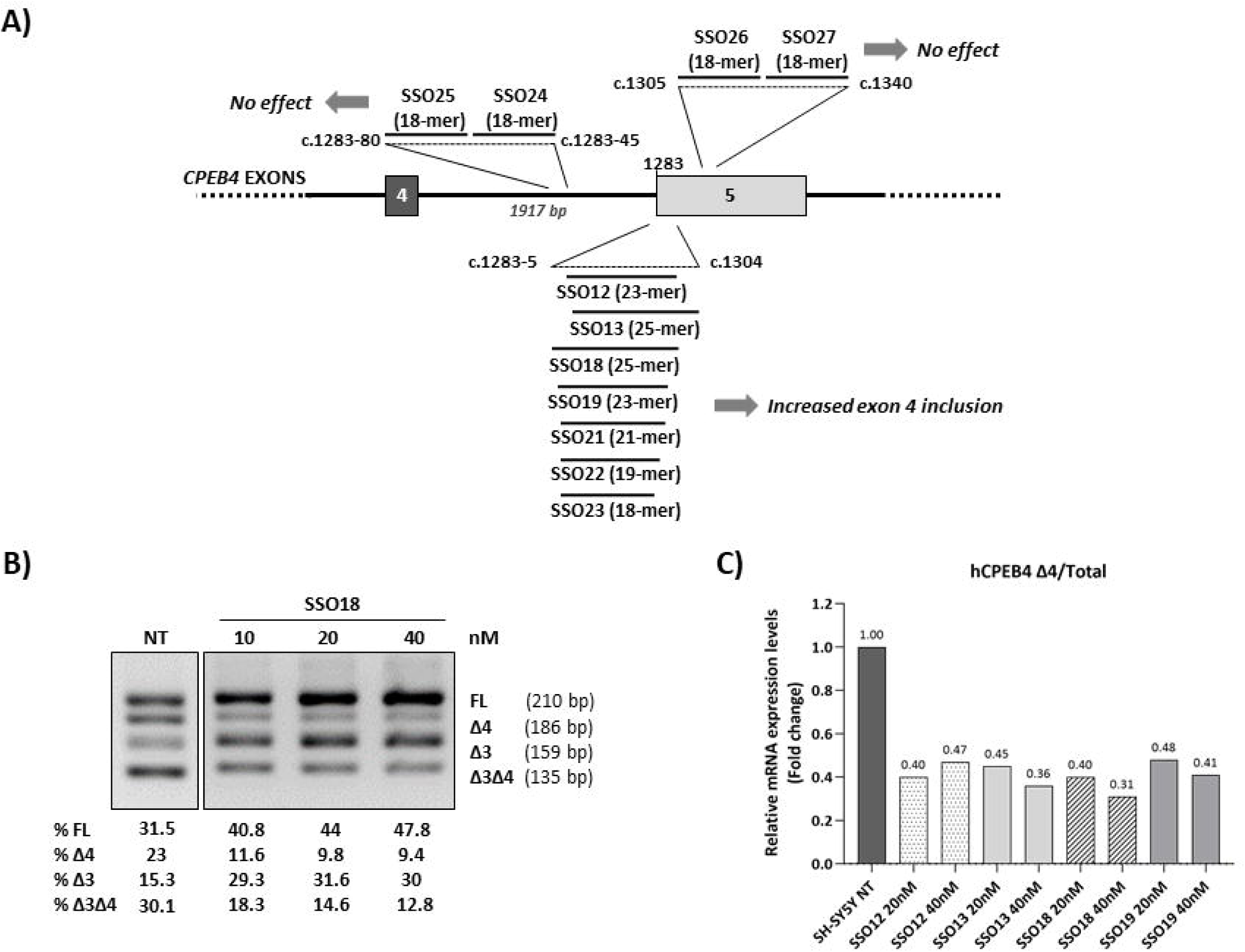
SSOs targeting the 3’ splice site of *CPEB4* exon 5 in SH-SY5Y cells. A) Schematics of the region with the location of the different SSOs and their effect on exon 4 inclusion. B) Gel showing the RT-PCR results after transfection with SSO18 at different concentrations. The identity of the bands is shown on the right and below is the estimation of the percentages of each band by laser densitometry. These results are representative of two independent experiments. C) RT-qPCR results after transfection with SSO12,13,18 and 19, expressed as fold change in the ratio of isoforms without exon 4 and total (includes all isoforms), relative to the levels in non-transfected cells (NT).

The initial results after transfection in SH-SY5Y with SSOs 12, 13, 18 and 19 (all of them 23 or 25-mer) showed that each of the SSOs stimulated inclusion of exon 4, decreasing the ratio CPEB4Δex4/total to 30-50% (Figure 3B, C). A dose-dependent effect was observed (Figure 3B, C).

Using SSOs with shorter lengths (18-, 19-, or 21-mer) targeting intron 4-exon 5 junction we observed a lower efficiency in decreasing Δex4 isoforms compared to the 23-mer SSO19 taken as reference (Supplementary Figure 1A). In addition, SSOs spanning upstream and downstream in the region (Figure 3A) allowed us to define the limits of the region amenable to SSOs targeting for favoring microexon inclusion. Overall, oligonucleotides targeting the region within −5 to +22 nucleotides relative to the first nucleotide of exon 5 promoted exon 4 inclusion. Both SSOs no. 24-25, which target the region upstream of nucleotide c.1283-45 and SSOs no. 26-27 hybridizing downstream in exon 5 had no effect (Supplementary Figure 1B). The use of a reverse primer in exon 6 confirmed that the SSOs did not result in skipping of exon 5 (Supplementary Figure 1B).

The SSOs that in SH-SY5Ycells showed a positive effect were also tested in murine N2A cells. We were able to recapitulate a similar effect (Supplementary Figure 2A-B), even though these SSOs were designed to target human sequences, and therefore, contain one or two mismatches with respect to the murine sequences. When N2A cells were transfected with equivalent SSOs designed with perfect complementarity to the murine sequences, larger decreases in Δ4 isoforms were detected (Supplementary Figure 2C-D).

### Minigene analyses point to the mechanisms underlying SSO effects

With the aim of investigating the mechanism underlying the pathogenic effect of SSO5 and the therapeutic effect of SSO19, which hinder or favour *CPEB4* microexon inclusion, respectively, we employed minigenes and targeted mutagenesis. A minigene with the human sequence of the *CPEB4* gene spanning from exon 2 to exon 6, with shortened introns, was cloned in the pcDNA3.1 vector (Figure 4A). Transfection in SH-SY5Y cells resulted in two transcripts, with (full length, FL) and without microexon 4 (Δ4) (Figure 4B-C). Despite testing different intron lengths in the minigene, exon 3 was not alternatively spliced in this system. Transfection of SSO5 and SSO19 reproduced the effects observed previously on microexon processing of the endogenous gene, by either promoting its exclusion (SSO5, increased Δ4 isoform) or its inclusion (SSO19, increased FL isoform) (Figure 4B, C), confirming the utility of the model.

**Figure 4.**
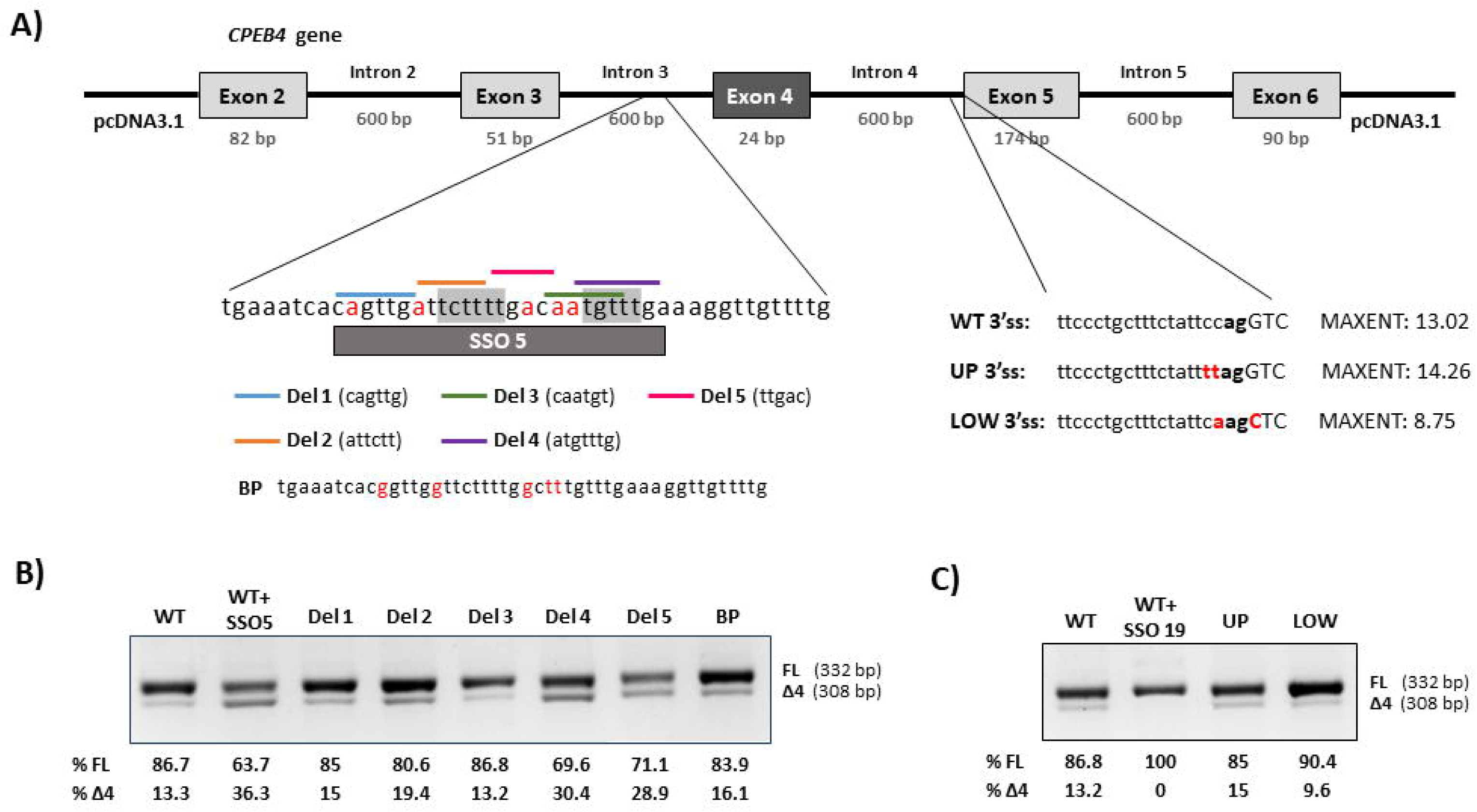
Minigenes analyses. A) Schematics of the *CPEB4* minigene construct and the deletions and point mutations introduced by mutagenesis. BP indicates mutations abolishing potential branch point sites. Mutations depicted as UP or LOW increase or decrease the 3’ splice site strength of exon 5, respectively, as shown by the resulting MaxEnt score (http://hollywood.mit.edu/burgelab/maxent/Xmaxentscan_scoreseq_acc.html) (B) RT-PCR results after transfection of mutant minigenes carrying deletions in intron 3 region, as compared to treatment of the wild-type (WT) minigene cotransfected or not with SSO5. C) RT-PCR results after transfection of mutant minigenes carrying point mutations in the 3’ splice acceptor site of exon 5, as compared to treatment of the WT minigene cotransfected or not with SSO19. These results are representative of two independent experiments. On the right of each gel is the schematic drawing showing the identity of the bands, determined by Sanger sequencing.

*CPEB4* microexon processing has been shown to be affected, at least in part, by manipulating SRRM4 levels [32]. SRRM4 forms a complex with SRSF11 and RNPS1, with SRSF11 recognizing an UCUCU motif [18]. Examination of the upstream intronic sequence of *CPEB4* microexon revealed two UCUUU(U) motifs in the region targeted by SSO5 (Figure 4A). In this region there are also predicted hnRNPC and hnRNPM motifs and potential BP sites (see Figure 2A) which could be responsible for the microexon exclusion observed with SSO5 treatment. To investigate the mechanism involved in the recognition of microexon 4 disrupted by SSO5 treatment, we introduced a series of deletions in the minigene covering the SSO5 target region. We also generated a mutant minigene in which all the A’s which could form part of potential BPs were changed to G or T nucleotides (Figure 4A). After transfection of the mutant minigenes in SH-SY5Y cells, followed by RT-PCR analysis, we observed that microexon 4 exclusion (Δ4 isoform) increased in the minigenes in which UCUUU(U) motifs are partially deleted (Del4, Del5 and, to a lesser extent, Del2), in a similar way as with SSO5 treatment (Figure 4B). With the other mutant minigenes, no significant effect on microexon inclusion was observed.

We also generated mutant minigenes with point mutations in the 3’ splice site of exon 5 that decreased or increased the splice site strength, as measured by the resulting MaxEnt splice score, to confirm if the effect of SSO19 was indeed related to the efficient recognition of the 3’ splice site downstream of the microexon. The results show that when the strength of the 3’ splice site of exon 5 is decreased, microexon inclusion is favoured, resembling what is observed with SSO19 treatment (Figure 4C).

### SSO5 binding blocks hnRNPC and PTPB1 binding sites

The mechanism of negative regulation of microexon splicing by SSO5 was further investigated using RNA affinity pull-down and subsequent Western blot analysis, in an attempt to identify RBPs whose binding to this region may be disrupted by competition with SSO5. We used biotinylated oligonucleotides covering the region targeted by SSO5 and available antibodies against SRRM4, RBFOX2, hnRNPC and PTPB1. Our results provide no evidence of binding of SRRM4 to this region and demonstrate specific binding of hnRNPC and PTPB1 (Figure 5).

**Figure 5.**
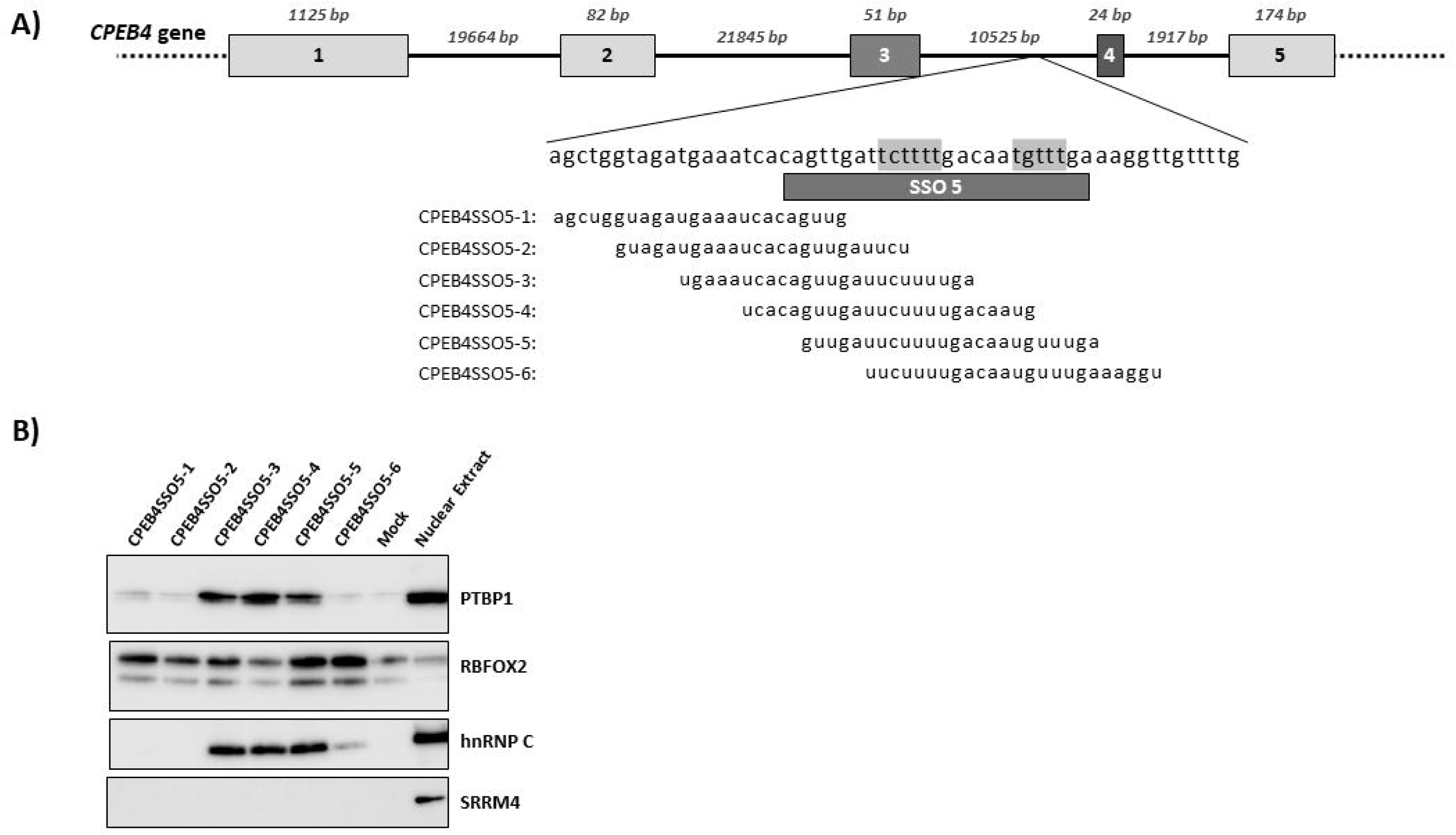
RNA affinity purification. A) Sequences of biotin-conjugated RNA oligonucleotides covering the SSO5 binding site in *CPEB4* intron 3. B) Western blot of SRRM4, RBFOX2, PTBP1 and hnRNPC proteins purified by RNA-affinity chromatography. Results are representative of two independent experiments. Mock and Nuclear Extract indicate control lanes without RNA oligonucleotides or with nuclear extract alone, respectively.

### *CPEB4* splicing modulation affects ASD-risk genes protein levels

We have previously shown that imbalance in *CPEB4* transcript isoforms that result from decreased inclusion of microexon 4 are associated with a decrease in poly(A)-tail length in high-confidence ASD risk genes, correlating with reduced expression of the protein products of ASD risk genes whose transcripts are binders of CPEB4 [6]. Therefore, to confirm a functional effect of the therapeutic value of the hit SSOs we tested whether SSOs promoting exon 4 inclusion affect protein levels of selected CPEB4-target ASD risk genes. To this aim we transfected N2A cells with mSSO18 (with mouse specific sequence) and examined by Western blotting levels of FOXP1 and AUTS2, protein products of *CPEB4* targets that exhibited decreased levels in TgCPEB4Δ4 mice [6]. For this, we previously confirmed that a murine specific version of SSO5 (mSSO5) promoted exon 4 skipping (emulating human specific SSO5 in SH-SY5Y cells), an effect that was counteracted by cotransfection of mSSO18. Both AUTS2 and FOXP1 levels were increased with mSSO18 treatment (Figure 6B). Of note, AUTS2 levels tended to decrease with mSSO5, while cotransfection of both mSSO5 and mSSO18 increased both AUTS2 and FOXP1 levels (Figure 6B).

**Figure 6.**
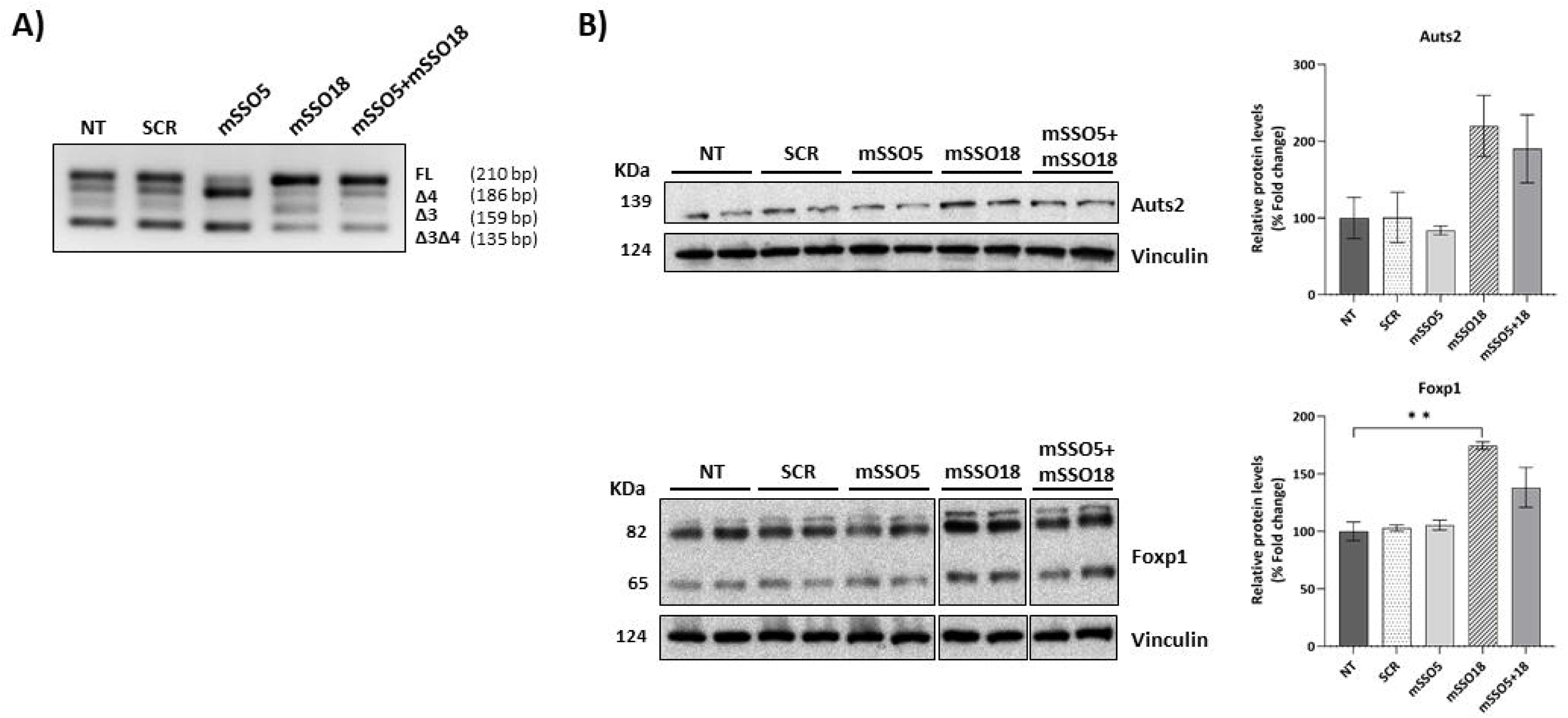
Analysis of *CPEB4* target genes protein levels after SSO transfection. (A) *CPEB4* RT-PCR analysis and (B) Western blot analysis of FOXP1 and AUTS2 levels in N2A cells after transfection with mSSO5 and mSSO18. Shown on the right of each gel are the relative amounts of FOXP1 and AUTS2 levels estimated by laser densitometry. Vinculin was used as loading control. NT: non-transfected; SCR: scrambled. Results are representative of three independent experiments Statistical analysis performed by one-way ANOVA (\*\**P*<0.01). Data are presented as mean ± SEM.

### Splicing of other neuronal microexons can be similarly modulated

We next considered whether targeting the downstream splice acceptor site could be a universal strategy to favor microexon inclusion, so we tested the same strategy for *TAF1* and *eIF4G1* microexons, which are both misregulated in brain diseases [32,33]. We designed SSOs hybridizing to the region encompassing the 3’ splice acceptor site downstream of microexon 34’ of the *TAF1* gene and of microexon 3 of the *eIF4G1* gene and tested their effect by RT-PCR after transfection in SH-SY5Y cells. Both SSOs hybridizing to the corresponding target region in the *TAF1* gene promoted microexon inclusion (Figure 7A), while no change was observed for *eIF4G1* microexon with the tested SSOs (Figure 7B).

**Figure 7.**
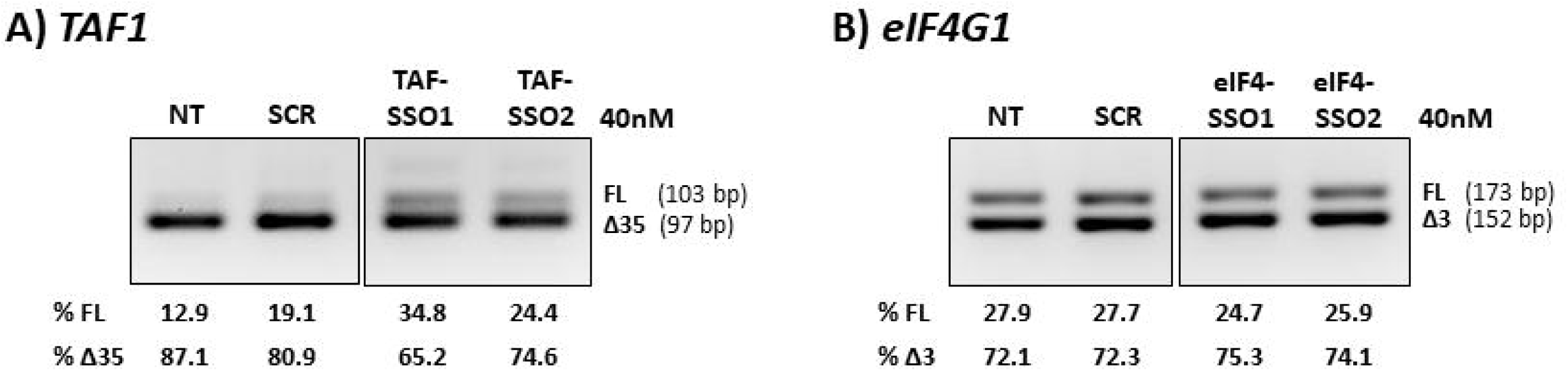
Effect of SSOs targeting the 3’ splice site downstream of the neuronal microexons in the *TAF1* (A) and *eIF4G1* (B) genes. The figure shows the gel analysis of RT-PCR after transfection with SSOs in SH-SY5Y cells. The identity of the bands is shown on the right and below is the estimation of the percentages of each band by laser densitometry. These results are representative of two independent experiments. NT: non-transfected; SCR: scrambled.

## DISCUSSION

AS affects more than 90% of human genes, with the brain characteristically exhibiting more AS events than any other tissue, indicative of its crucial role in neural development and function [34]. Indeed, AS dysregulation has emerged as a frequent mechanism involved in the pathogenesis of neurodegenerative diseases such as Alzheimer’s disease, Parkinson’s disease, Huntington’s disease, amyotrophic lateral sclerosis and frontotemporal dementia [34,35]. Furthermore, evidence has accumulated of the presence of neuronal specific microexons undergoing AS, which constitute a regulatory network that is disrupted in ASD [3]. Several of these neuronal microexons that impact critical neuronal processes, and whose mis-splicing results in distinctive autistic features, have been characterized to date [5], among them the 24 bp *CPEB4* microexon 4, which is the main focus of this study. Disruption of *CPEB4* neuronal microexon splicing, leading to an imbalance in transcript isoforms, has emerged as a common mechanism underlying both ASD and SCZ [6,14].

Upon identification of mis-spliced neuronal microexons with critical roles in ASD and other neurological diseases, the next challenge is to test existing therapeutic approaches known to modulate splicing, such as SSOs, as a way to revert specific disease phenotypes. In this work we sought to identify SSOs which could promote microexon inclusion in neuroblastoma cells, of potential use to correct the isoform imbalance associated with ASD and SCZ.

Our first strategy consisted in targeting intronic regions flanking the microexon. This first screen identified SSO5 that, contrary to what we aimed for, promoted microexon exclusion, thus mimicking a pathogenic state. Interestingly, no cellular model of *CPEB4* mis-splicing as occurring in ASD or SCZ is available to date, so this finding represents a way of generating a useful cellular tool for the analysis of related pathogenic mechanisms in these neuropsychiatric diseases, as well as to test therapeutic approaches. The approach is reminiscent of the method named TSUNAMI (targeting splicing using negative ASOs to model illness), whereby Krainer and coworkers used SSOs to induce *SMN2* mis-splicing *in vitro* and *in vivo*, thus accurately phenocopying spinal muscular atrophy [36,37].

The mechanism of SSO5-mediated microexon exclusion is most probably related to competition with sequence specific RBPs binding to the region. RNA pulldown experiments indicate no binding of SRRM4 to this region, thus ruling out the involvement of the complex including SRRM4/SRSF11 previously described to regulate neuronal microexons [32]. Our results point to other RBPs that have also been reported to participate in microexon usage such as RBFOX and PTBP1[38], as SSO5 competes with hnRNPC (and indirectly also with RBFOX1, as it coexists with hnRNPC, hnRNP M, hnRNP H and other proteins in the LASR complex [39]), as well as with PTPB1, which binds to a splicing enhancer region containing two U-rich motifs, upstream of the microexon, thus promoting its exclusion. This effect was reproduced when these motifs were deleted using minigenes. One limitation of the minigene model is that it did not fully recapitulate the transcript profile of the *CPEB4* gene, as no AS of exon 3 could be detected, probably due to lack of the complete genomic context.

The main result of our work is the identification of SSOs targeting the 3’ splice acceptor site sequence downstream of the microexon, which effectively promote its incorporation in the mature transcript. To our knowledge, this is the first time that SSO-mediated microexon inclusion has been achieved. Employing SSOs with different lengths we could show that stimulation of microexon 4 splicing was dependent on the strength of SSO hybridization to the 3’ splice site, with shorter SSOs resulting in lower levels of microexon inclusion. SSOs predictably compete with spliceosomal components binding to the same site, thus slowing down spliceosomal assembly and transcript elongation and lengthening the window of opportunity for upstream microexon recognition and processing. This is in accordance to the kinetic coupling model model that proposes that changes in transcriptional elongation rates can regulate splicing by modulating the presence of competing splicing sites[20].

This mechanism may also operate for other AS microexons, as shown in this work for the 6 bp microexon of *TAF1*. However, this splice modulating approach does not operate for all microexons, as seen for the *eIF4G1* microexon. The different results may be related to the splicing strength, estimated as the calculated splice score, of the targeted downstream 3’ splice site (MaxEnt scores: for *TAF1* it is 7,22; for *CPEB4 it is* 7,67 and for *eIF4G* 8,22), making the competition of SSOs with the spliceosomal components more difficult. In addition, it is known that both the genomic and the epigenetic context may act modulating the transcription rate and thus the associated splicing process of each exon [20]. More studies are necessary to understand the gene and exon-specific features that influence the splicing outcome of this approach.

The functional effect of this innovative SSO-mediated modulation of the ratio of *CPEB4* transcript isoforms was shown by the increase in the levels of FOXP1 and AUTS2 proteins, products of ASD risk genes, whose transcripts are binders of CPEB4. This provides proof-of-concept of the potential therapeutic effect of the SSO, by correcting the concerted mis-expression of the network of high-confidence ASD risk genes regulated by *CPEB4,* identified in brains of idiopathic ASD cases and in SCZ [6,14].

Only a small fraction of ASD cases are monogenic and caused by penetrant mutations. Several corrective therapies for ASD are in preclinical/clinical development, such as gene therapy [40] or oligonucleotides targeting pseudoexons/non-productive AS events [41–43]. These therapies are specific for a given gene/mutation and thus are applicable to only some of those genetic forms of ASD. However, we have previously shown that *CPEB4* mis-splicing occurs in the more prevalent forms known as idiopathic ASD, driving concerted mis-expression of a plethora of high confidence ASD risk genes in the brain of idiopathic ASD cases [6]. Thus, our results shown here open the opportunity to broaden the application of SSOs to idiopathic forms, and potentially also to certain genetic forms of ASD in which convergent mechanisms of under-expression of neurodevelopmental genes may be operating.

Antisense therapeutic approaches leverage on the continuous and ongoing improvements in oligonucleotide chemical modifications to enhance stability and potency and in innovative delivery methods to ensure a successful transition from bench to bedside. SSOs are in clinical use for Duchenne muscular dystrophy and spinal muscular atrophy, and individualized treatments approved for clinical testing in n=1 trials are emerging [44,45]. A number of SSO are also in clinical development for Huntington’s disease, amyotrophic lateral sclerosis, Parkinson’s disease and Alzheimer’s disease [34]. Results as those presented here highlight the potential of RNA splicing therapies for previously intractable diseases.

## DECLARATIONS

### Data availability statement

All of the relevant data supporting the findings of this study are presented in the article or in the supplemental material. Additional information and other raw data are available from the corresponding authors upon reasonable request.

### Authors contributions

JJL, BSA and LRD conceptualized this study and provided supervision. AMP and SP performed the SSO transfections, RT-PCR and western blotting experiments. LLH and TKD performed RNA affinity pulldowns. AMP, JJL, BSA and LRD drafted and revised the manuscript. All of the authors reviewed the final draft.

## Acknowledgements

The technical support of Mar Alvarez and the help of Beatriz Alvera is gratefully acknowledged. We thank Julia Pose-Utrilla for the critical reading of the manuscript and useful suggestions.

