## Supplementary material for "OLIGONUCLEOTIDES TARGETING THE 3’ SPLICE SITE DOWNSTREAM OF A MICROEXON AS AN INNOVATIVE THERAPY FOR AUTISM"

### *CPEB4* MIS-SPLICING CORRECTION WITH ANTISENSE OLIGONUCLEOTIDES AS POTENTIAL THERAPEUTIC STRATEGY FOR AUTISM AND OTHER BRAIN DISORDERS

Ainhoa Martinez-Pizarro, Sara Picó, Lise Lolle Holm, Thomas K Doktor, Brage S Andresen, José J Lucas, Lourdes R Desviat

**Supplementary Table 1. SSO sequences**

| Gene | Name | Sequence (5′-3′) | Mer |
|---|---|---|---|
| **Human SSOs** | | | |
| | **Scrambled** | CUCAAUAUGCUACUGCCAUG | 20 |
| *CPEB4* | **SSO1** | AGUGAUACAUUUAAAAGUUAAAAAC | 25 |
| | **SSO2** | CAAGUGAUACAUUUAAAAGUUAAAA | 25 |
| | **SSO3** | UACAAGUGAUACAUUUAAAAGUUAA | 25 |
| | **SSO5** | CAAACAUUGUCAAAAGAAUCAACUG | 25 |
| | **SSO6** | AAACAAAAAAACAAAACAACCUUUC | 25 |
| | **SSO7** | CUCCCCUUCCCCACAAAGGUUAGAA | 25 |
| | **SSO8** | AAAUUACCUAAGGAUCAAAGCUAUG | 25 |
| | **SSO12** | UGGAAACAGUGAAGACUGACCUG | 23 |
| | **SSO13** | AUUGGAAACAGUGAAGACUGACCUG | 25 |
| | **SSO18** | UGGAAACAGUGAAGACUGACCUGGA | 25 |
| | **SSO19** | GAAACAGUGAAGACUGACCUGGA | 23 |
| | **SSO20** | GGAAACAGUGAAGACUGACCUGG | 23 |
| | **SSO21** | AAACAGUGAAGACUGACCUGG | 21 |
| | **SSO22** | ACAGUGAAGACUGACCUGG | 19 |
| | **SSO23** | CAGUGAAGACUGACCUGG | 18 |
| | **SSO24** | UGCCACUGAUUAUCAUGC | 18 |
| | **SSO25** | AAAGAAACAAGGAAACAG | 18 |
| | **SSO26** | UCCAAGAAUCCAUCUUCC | 18 |
| | **SSO27** | UGAUCCCCACGGCCAUCA | 18 |
| *TAF1* | **SSO1** | GUAUCAUACAAAUCAGGAGGCUGUG | 25 |
| | **SSO2** | AUCAUACAAAUCAGGAGGCUGUG | 23 |
| *eIF4G1* | **SSO1** | GCCCGGCUAGGGUAGAAGUGCUGUC | 25 |
| | **SSO2** | CCGGCUAGGGUAGAAGUGCUGUC | 23 |
| **Mouse SSOs** | | | |
| *Cpeb4* | **mSSO5** | CAAACAUUGUCAAAGAAUCCACUG | 24 |
| | **mSSO12** | CGGAAACAAUGAAGACUGACCUG | 23 |
| | **mSSO13** | AUCGGAAACAAUGAAGACUGACCUG | 25 |
| | **mSSO18** | CGGAAACAAUGAAGACUGACCUGGA | 25 |
| | **mSSO19** | GAAACAAUGAAGACUGACCUGGA | 23 |

**Supplementary Table 2. Primer sequences**

| Tecnique | Gene | Name | Sequence (5′-3′) |
|---|---|---|---|
| PCR | *CPEB4* | **mhExon 2 F** | GGACGTTTGACATGCACTCAC |
| | | **hExon 3 F** | ATCCAGGATCCGATAGCTCTC |
| | | **mhExon 5 R** | GAGGTTGATCCCCACGGC |
| | | **hExon 6 R** | CCAAAGCGACGAAAACTAGC |
| | | **hExon 6-pcDNA3.1 R** | CTAGACTCGAGCTTTAGGAGG |
| | *TAF1* | **hExon 34 F** | TAGAAAGCCTGGACCCAATG |
| | | **hExon 36 R** | GAGGCATCTCGAGACATACTGA |
| | *eIF4G1* | **hExon 2 F** | TCAGTACGCCACAAGCGAC |
| | | **hExon 4 R** | AGCAGGGTAGACATGGGCAG |
| qPCR | *CPEB4* | **hCPEB4 Δ4 F** | ATCCGATAGCTCTCTGCTTATTAATGGT |
| | | **hCPEB4 Δ4 R** | CACGGCCATCATCCAAGAAT |
| | | **hCPEB4 total F** | CACTGTTTCCAATGGAAGATGG |
| | | **hCPEB4 total R** | GGTGAACCCAGGCCACTATG |
| | *GAPDH* | **hGAPDH F** | GTCGGAGTCAACGGATTTGG |
| | | **hGAPDH R** | GACAAGCTTCCCGTTCTCAG |
| | *Cpeb4* | **mCpeb4 4 F** | ATCCGATAGTTCTCTGCTTATTAATGGT |
| | | **mCpeb4 4 R** | ACGGCCATCATCCAGGAAT |
| | | **mCpeb4 total F** | CAAATCTTATTTTCCACCAAAAGG |
| | | **mCpeb4 total R** | CATCAATGAGAGCCTGAACAGA |
| | *Gapdh* | **mGapdh F** | AGCTGAACGGGAAGCTCACT |
| | | **mGapdh R** | GCTTCACCACCTTCTTGATGTC |

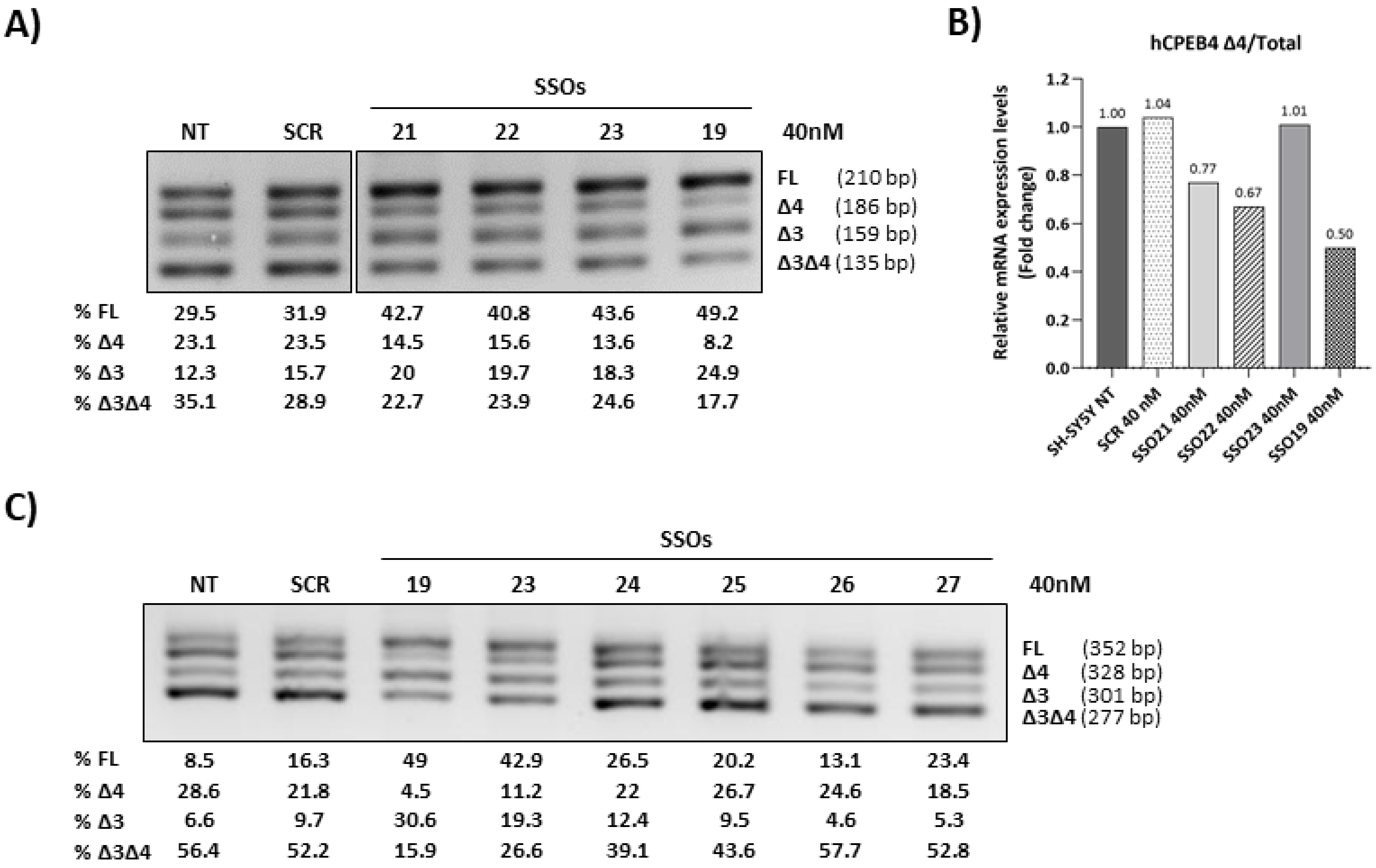


**Supplementary Figure 1. SSOs targeting the 3' splice site of CPEB4 exon 5.** (A) RT-PCR results after transfection in SH-SY5Y cells with SSOs of different lengths at 40 nM. (B) quantitative RT-qPCR results expressed as fold change in the ratio of isoforms without exon 4 and total (includes all isoforms), relative to the levels in non-transfected cells. (C) RT-PCR results using a reverse primer in exon 6 after transfection in SH-SY5Y cells with SSOs 24-27 targeting upstream and in exon 5, with SSO19 and SSO23 taken as reference. The identity of the bands is shown on the right and below is the estimation of the percentages of each band by laser densitometry. The results are representative of two independent experiments. NT: non-transfected. SCR: scrambled SSO.

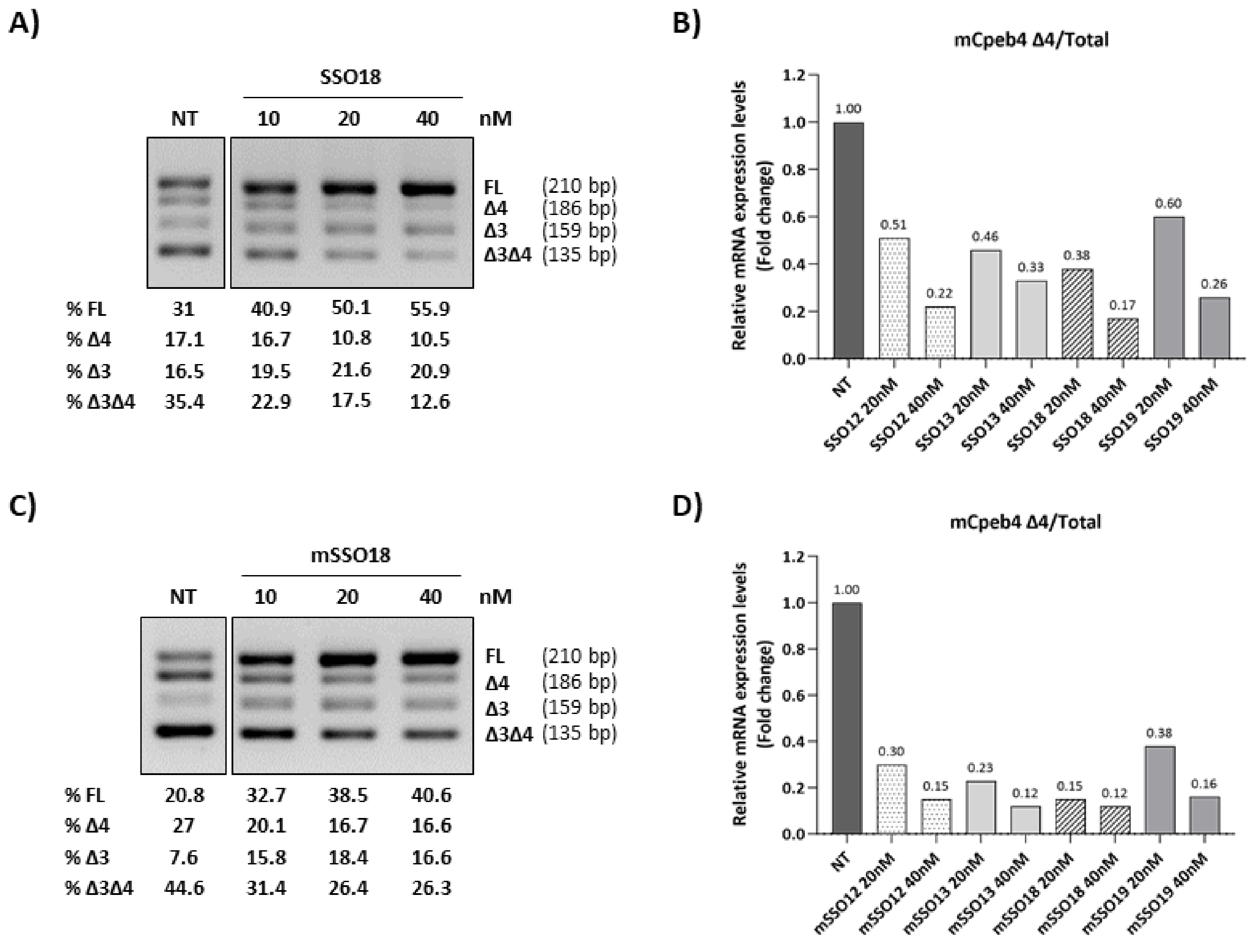


**Supplementary Figure 2: SSOs targeting the 3' splice site of CPEB4 exon 5 favor microexon 4 inclusion in N2A cells.** (A) Gel showing the semiquantitative RT-PCR results after transfection with SSO18 at different concentrations. (B) RT-qPCR results expressed as fold change in the ratio of isoforms without exon 4 and total (includes all isoforms), relative to the levels in non-transfected cells (NT). (C) and (D) are the same as (A) and (B) using the corresponding murine sequences of each SSO. The identity of the bands is shown on the right and below is the estimation of the percentages of each band by laser densitometry.
